# Patterns of serum immune biomarkers during elephant endotheliotropic herpesvirus viremia in Asian and African elephants

**DOI:** 10.1101/2021.05.12.443748

**Authors:** Katie L. Edwards, Erin M. Latimer, Jessica Siegal-Willott, Wendy Kiso, Luis R. Padilla, Carlos R. Sanchez, Dennis Schmitt, Janine L. Brown

## Abstract

Hemorrhagic disease (HD) caused by a group of elephant endotheliotropic herpesviruses (EEHV) is one of the leading causes of death for young elephants in human care. These viruses are widespread and typically persist latently in adult elephants with no negative effects; however, in juvenile Asian and more recently young African elephants, the onset of disease can be rapid and the mortality rate high. Measuring biomarkers associated with the immune response could be beneficial to understanding underlying disease processes, as well as the management of infection and HD. The goal of this study was to measure acute phase proteins and cytokines in serum collected from elephants infected with EEHV (13 Asian and 1 African) and compare concentrations according to presence, severity and outcome of disease. Serum amyloid A (SAA) and haptoglobin (HP) were higher in elephants with EEHV viremia than those without; concentrations increased with increasing viral load, and were higher in fatal cases compared to those that survived. In Asian elephants, SAA was also higher during EEHV1 viremia compared to EEHV5. Cytokine concentrations were typically low, and no statistical differences existed between groups, perhaps due to insufficient assay sensitivity in this age group. However, in individuals with detectable levels, longitudinal profiles indicated changes in tumor necrosis factor alpha (TNF-α) and interleukin-2 (IL-2) that may reflect an immune response to EEHV infection. These results highlight the potential benefit of measuring circulating biomarker concentrations, such as APPs and cytokines, to improve our understanding of EEHV viremia and HD, assist with monitoring the progression of disease and determining the impact of interventions.

## Introduction

Over the past two decades, an acute hemorrhagic disease (HD) caused by a group of elephant endotheliotropic herpesviruses (EEHV) has contributed to 65% of deaths of young, captive-born Asian elephants in North American and European zoos [1], affecting almost one in four globally [2]. EEHV-HD primarily occurs in Asian elephants under 10 years of age, but has also contributed to the death of older individuals, as well as African elephants [3-6]. Once thought to affect only western *ex situ* collections, cases of EEHV HD have now been reported in captive and wild populations in India [1, 7-9], Thailand [10-14], Laos [15], Cambodia [16], Myanmar [17], Nepal, and Sumatra [18, 19].

Zoo studies demonstrate that many Asian and African elephants intermittently shed EEHV DNA via oronasal mucosa and perhaps also through ocular and urogenital secretions [20-25], with serological investigations finding widespread latent or subclinical infection [11, 18, 26]. Of the seven major sub-types, EEHV1, 4 and 5 are endemic to Asian elephants, whereas EEHV2, 3, 6, and 7 are found in African elephants [19]. Although low-level viremia may be fairly ubiquitous across both species [5, 27], in Asian elephant calves between 1-8 years of age, increasing viremia and the onset of EEHV HD can be rapid, resulting in death within a few hours to days as widespread endothelial cell necrosis occurs [28, 29]. Most of the deaths to-date have resulted from EEHV1A and EEHV1B infection in Asian elephants, with fewer fatalities associated with EEHV4 and 5. Recent deaths of young African elephants (6 to 11 years of age) have been associated with EEHV3A [4-6], with HD also associated with EEHV3B [30]. Early detection through routine testing and immediate intervention with supportive care and anti-viral therapies are critical to successful outcomes [18]. However, what remains unclear is why some individuals quickly succumb to EEHV HD while others that develop high viral loads survive [30-32]. Recent evidence suggests that severe cases of EEHV HD result from primary infection, as individuals that died were seronegative for EEHV-specific antibodies [33], and that T-cells likely are key to fighting EEHV infection [34]. Certainly, more detailed knowledge of how the host immune system responds to infection, and any differences between fatal and surviving cases of EEHV viremia and HD would be beneficial to understanding disease pathogenesis, the development of targeted treatments, and ultimately reducing morbidity and mortality from this disease.

The typical immune response to a viral infection includes both innate and adaptive components [35-37]. Cytokines play an important role in recruiting, activating and otherwise moderating immune cell function to facilitate the initiation and development of the immune response. They are involved in both innate and adaptive immunity, which results in the body developing pathogen-specific T-cells and antibodies, generally over the course of several days post-infection. Tumor necrosis factor (TNF)-α and interleukin (IL)-6 [38], interferon (IFN)-γ [39], interleukin-2 [40] and interleukin-10 [39] all have been associated with herpesvirus infections in other species, and expression of cytokine mRNA including TNF-α and IFN-γ is increased in Asian elephants persistently infected with EEHV4 [41]. Although circulating cytokine concentrations have not been previously investigated with respect to the development of EEHV HD specifically, they play a role in the elephant immune response to tuberculosis [42-44] and other infectious and non-infectious pathologies [45]. Acute phase proteins (APPs) form part of the innate acute phase response against infection or tissue injury and as such are among the first signs of inflammation. Two APPs, serum amyloid A (SAA) and haptoglobin (HP) [46], have previously been investigated in response to EEHV1 infection [29]. This study will add to that knowledge by analyzing serum samples from Asian and African elephants during EEHV viremia to compare concentrations of APPs SAA and HP, and cytokines TNF-α and IL-2, which are pro-inflammatory and associated with T-cell growth and differentiation, respectively. We investigated how these measures differ during viremia and non-viremia, among different types of EEHV, with increasing viral load, and between fatal and surviving cases of HD.

## Materials and methods

### Subjects and sample collection

Serum and whole blood samples submitted to the National Elephant Herpesvirus Laboratory (NEHL) at the National Zoological Park, Washington D.C. for routine EEHV testing were utilized in this study. Blood samples were collected according to approved phlebotomy protocols at each institution, typically from an ear vein through behavioral conditioning, or as part of a medical intervention. Whole blood was collected into EDTA anticoagulant tubes, refrigerated and shipped overnight to the NEHL. A second blood sample collected into serum tubes was allowed to clot at room temperature, before serum was separated, frozen at -20 to -80°C and shipped overnight to the NEHL or the Smithsonian Conservation Biology Institute. The presence of EEHV in whole blood was determined by conventional PCR (cPCR) and/or quantified by real-time PCR (qPCR) according to methodology already described by Latimer et al. [47], Stanton et al. [22] and Bauer et al. [48]. Viral load is presented as viral genome equivalent per milliliter (vge/ml).

This study included serum samples from 14 elephants (13 Asian and 1 African), aged 1 year 2 months to 12 years and 1 months at the time of collection (Table 1). For some individuals, whole blood and serum samples were submitted weekly to the NEHL for routine EEHV surveillance, allowing longitudinal analysis of biomarkers; in others, serum was submitted opportunistically. This research was approved by the management at each participating institution, and where applicable, was reviewed and approved by zoo research committees. The study protocol was also reviewed and approved by the Smithsonian’s National Zoo and Conservation Biology Institute (NZP-ACUC #15-03 and #18-18).

**Table 1:** Subjects and serum samples included in the study. The age range incorporates the period of viremia detection and may encompass multiple episodes over the study period.

| Animal ID# | Species | Sex | Age (years, months) | Number of viremia episodes<br>(by EEHV type) | No. serum<br>samples |
| --- | --- | --- | --- | --- | --- |
| 1 | *E.m* | F | 4 y 7 m - 5 y 3 m | 11 (5 EEHV1; 6 EEHV5) | 73 |
| 2 | *E.m* | F | 7 y 2 m - 7 y 7 m | 2 (1 EEHV1; 1 EEHV5) | 43 |
| 3 | *E.m* | F | 11 y 0 m - 11 y 4 m | 3 (EEHV5) | 40 |
| 4 | *E.m* | M | 3 y 8 m - 3 y 10 m | 1 (EEHV1) | 8 |
| 5 | *E.m* | F | 1 y 11 m - 11 y 8 m | 6 (3 EEHV1; 3 EEHV5) | 19 |
| 6 | *E.m* | F | 3 y 6 m - 12 y 0 m | 4 (1 EEHV1; 3 EEHV5) | 27 |
| 7 | *E.m* | F | 1 y 1 m - 5 y 5 m | 5 (2 EEHV1; 3 EEHV5) | 40 |
| 8 | *E.m* | F | 2 y 10 m - 6 y 7 m | 4 (1 EEHV1; 3 EEHV5) | 21 |
| 9 | *E.m* | F | 4 y 3 m - 4 y 5 m | 1 (EEHV1) | 4 |
| 10 | *E.m* | F | 1 y 3 m - 1 y 9 m | 2 (EEHV1) | 7 |
| 11 | *E.m* | M | 2 y 8 m - 5 y 11 m | 2 (1 EEHV1, EEHV 5) | 36 |
| 12 | *E.m* | F | 1 y 9 m - 5 y 11 m | 2 (1 EEHV1; 1 EEHV5) | 3 |
| 13 | *E.m* | F | 1 y 7 m - 5 y 8 m | 1 (EEHV1) | 4 |
| 14 | *L.a* | M | 4 y 11 m - 5 y 0 m | 1 (EEHV3B) | 28 |

### Serum biomarker quantification

SAA and HP concentrations were measured using a RX Daytona automated clinical chemistry analyzer (Randox Industries-US Ltd., Kearneysville, WV, USA). Commercially available reagents, calibrators, and two-level controls were used for each assay (Eiken Chemical Co. Ltd, Tokyo, Japan and Tridelta Tri-DD, Boonton, NJ, USA, respectively), which have previously been validated for elephants [29, 45, 46]. The technical ranges were 0.1-500 mg/l and 0.01-2.5 mg/ml, respectively. The analyzer was subject to routine quality control measurements throughout the study, with normal and elevated controls for each analyte maintained within two standard deviations (SD) of the respective manufacturer-defined target value. Samples were typically run neat, but those with HP above the technical range were diluted 1:5 in calibrator diluent.

Cytokines were measured by enzyme immunoassay as described in Edwards et al. [45]. In brief, TNF-α was measured using a commercially available equine TNF-α ELISA reagent kit (ESS0017; Thermo Fisher Scientific, Frederick, MD, USA), with anti-equine TNF-α coating antibody and biotinylated anti-equine TNF-α detection antibody both diluted 1:100, and recombinant equine TNF-α standards (3.9 to 1000 pg/ml). Interleukin 2 (IL-2) was measured using a commercially available equine Duoset ELISA development kit (DY1613) with modifications, incorporating anti-equine IL-2 coating (2.0 μg/ml) and biotinylated detection (0.2 μg/ml) antibodies and recombinant equine IL-2 standards (15.6 to 4000 pg/ml). Preliminary testing of samples from three individuals on other cytokine assays that have been validated for elephants (IFN-γ, IL-6, IL-10; [45]) were all below assay detection limits, and so were excluded from the remainder of the study. High and low controls within the standard range for each assay were maintained within an inter-assay CV of 15%. Serum was analyzed undiluted.

### Statistical analyses

Serum samples were categorized as viremia positive or negative, by EEHV type, and for viral load based on qPCR testing of whole blood collected on the same day. Data were then analyzed using generalized linear mixed models constructed using the package ‘lmer’ [49] in R [50], using a Gamma distribution with a log-link to model biomarkers concentrations. Due to longitudinal sampling, sample date and animal ID were included as random effects; viremia (positive or negative), EEHV type (EEHV1 or EEHV5) and viral load (0, 10^2^, 10^3^, 10^4^, 10^5^ and 10^6^ or above) were included as categorical fixed effects. Samples from the one African elephant with EEHV3B were excluded from the comparison of EEHV type. In addition, including only samples from cases of higher EEHV viremia (> 5000 vge/ml), concentrations of each biomarker were compared between cases that resulted in either survival or fatality. Animal ID and EEHV type were included as random effects, and data were modelled either using all samples between the start and end of the case of viremia, or using the peak concentration for each biomarker. All results are presented as the mean prediction ± standard error from the GLMM to account for repeated sampling over time within individuals.

## Results

Of 353 serum samples analyzed, 233 were categorized as positive by either cPCR or qPCR. These contributed to 46 separate cases of viremia, six of which were categorized as positive through cPCR but viral load was not quantified, 17 were transient low concentrations (< 1000 vge/ml), six were intermediate (1000-5000 vge/ml), and 17 were higher level viremia (> 5000 vge/ml; Table 2). Of these 46 cases, 20 were EEHV1, 25 were EEHV5 (all in Asian elephants), and the one African case was EEHV3B.

**Table 2:**
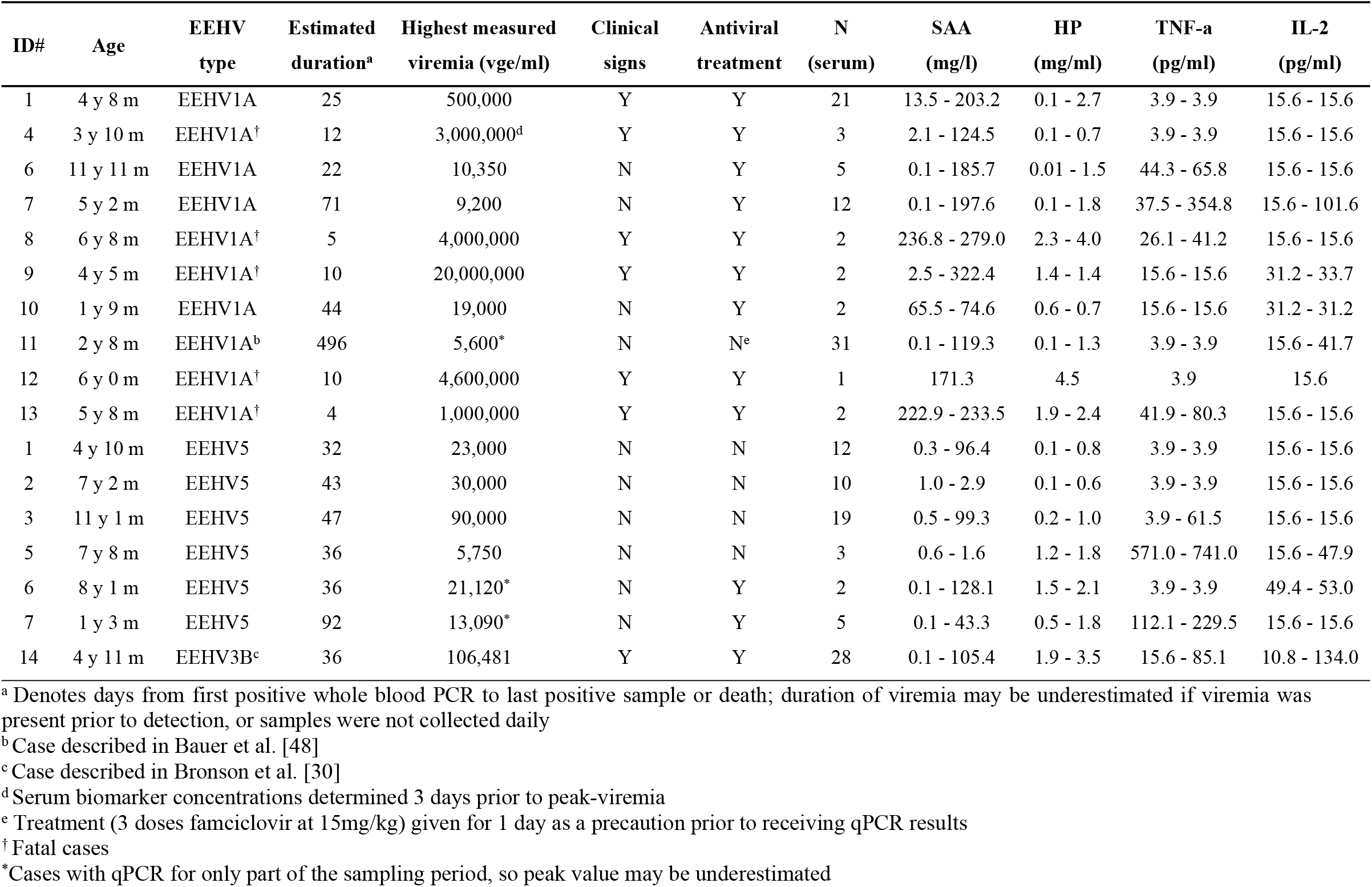
Range (minimum-maximum) of serum biomarker concentrations during cases of viremia exceeding 5000 vge/ml whole blood in 1 African and 13 Asian elephants.

Both SAA (P < 0.001) and HP (P = 0.009) were increased in EEHV-positive samples (38.1 ± 4.4 mg/l and 1.0 ± 0.1 mg/ml, respectively) compared to those that did not contain detectable EEHV (14.8 ± 4.0 mg/l and 0.6 ± 0.1 mg/ml, respectively); TNF-α and IL-2 did not differ between positive and negative samples (Fig 1). SAA was also higher (P = 0.002) in EEHV1 (55.2 ± 7.2 mg/l) viremia compared to EEHV5 (19.9 ± 4.9 mg/l); the other biomarkers did not differ by EEHV type in Asian elephants (Fig 2). Both APPs increased with increasing viral load (Fig 3); SAA was elevated in samples with viral load at 10^3^ vge/ml and above, compared to 0 vge/ml, and HP was elevated in samples with a viral load of 10^4^ and 10^5^ compared to 0 vge/ml. TNF-α did not differ significantly with increasing viral load; IL-2 concentrations were lower at 10^6^ vge/ml compared to 0 vge/ml (P = 0.042), although this relationship was driven by a single elevated IL-2 value in an Asian elephant calf with a swollen front limb that was not associated with EEHV viremia. Of the 17 cases of higher-level viremia (i.e. those that included peak viral load of > 5000 vge/ml; Table 2), an observable increase in SAA was detected in 15 cases, HP in 14 cases, TNF-α in five cases and IL-2 in two cases. Of the elephants with high viral loads, overall concentrations of SAA were higher in fatal cases of EEHV HD (P < 0.001), with a tendency for higher concentrations of HP (P = 0.088). Furthermore, peak concentrations were higher for HP (P < 0.001), and lower for IL-2 (P < 0.001) in fatal cases of EEHV HD compared to those that survived, with a tendency for higher peak SAA (P = 0.063).

**Fig 1:**
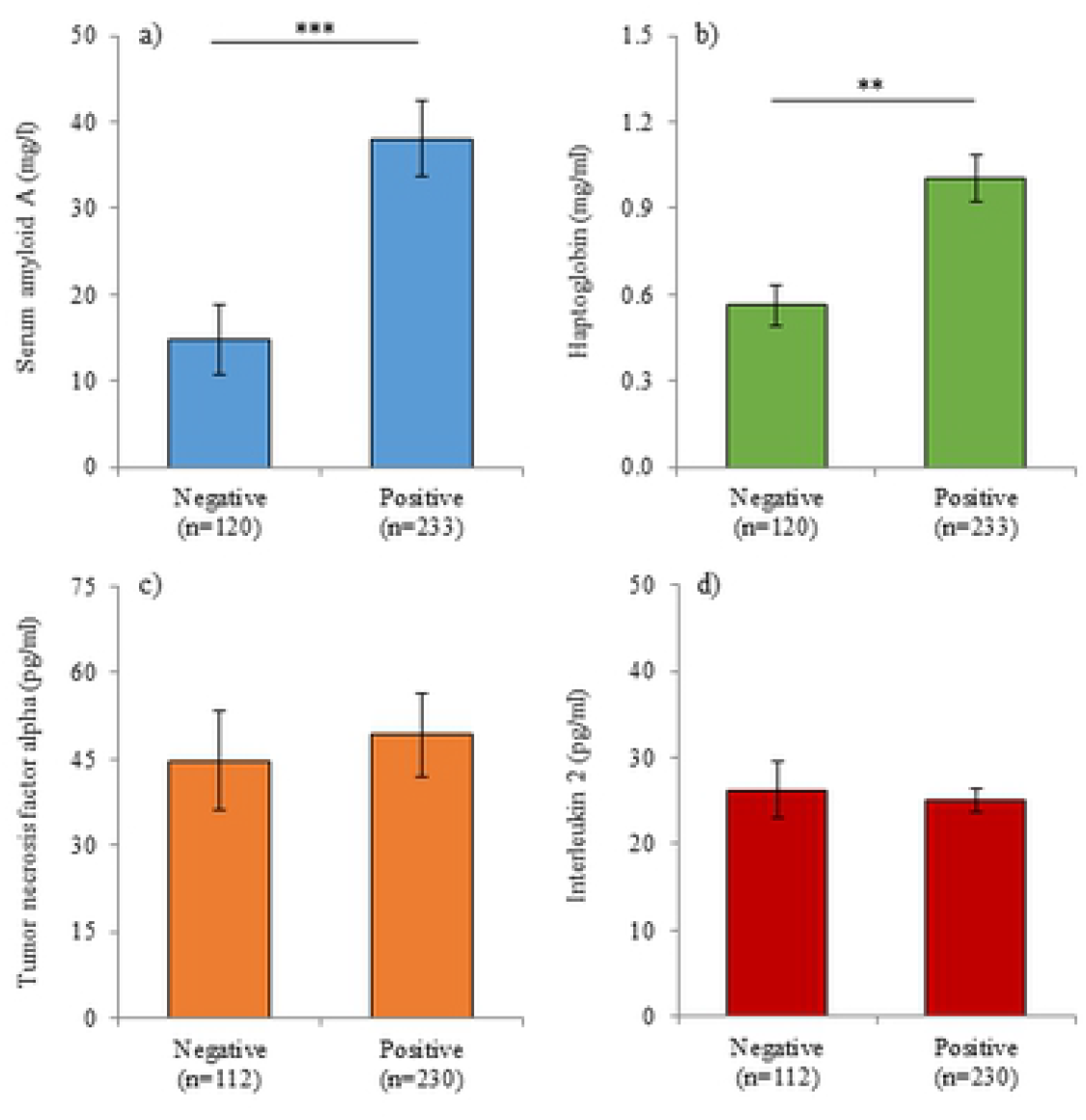
Acute phase protein and cytokine concentrations with positive or negative viremia. Serum concentrations of serum amyloid A (a), haptoglobin (b), tumor necrosis factor alpha (c) and interleukin 2 (d) on days when whole blood was positive or negative for EEHV viremia by PCR. Bars represent the mean ± sem of the prediction from the GLMM; asterisks denote significant differences between categories (** P < 0.01; *** P < 0.001).

**Fig 2:**
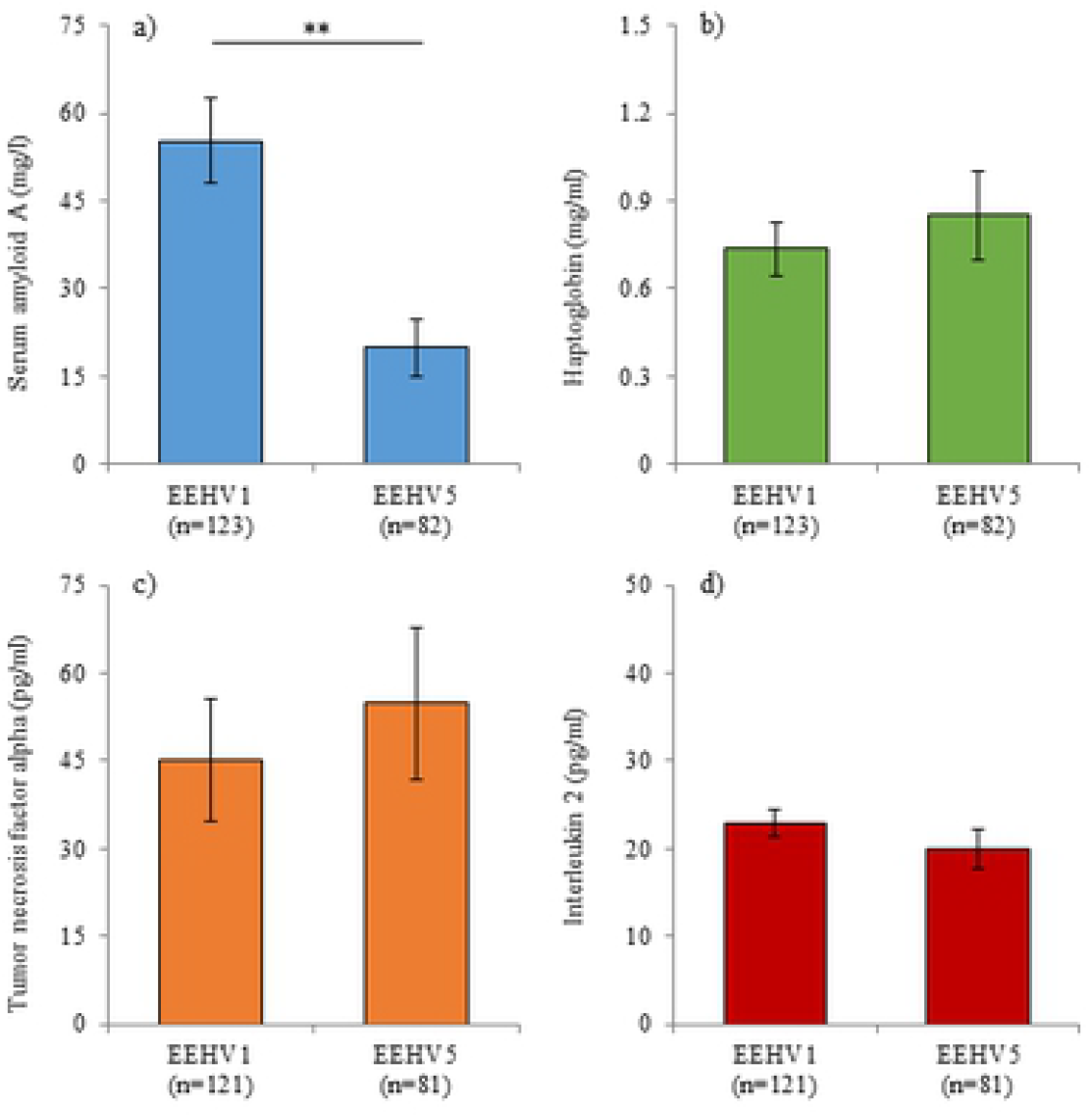
Acute phase protein and cytokine concentrations with EEHV1 or EEHV5. Serum concentrations of serum amyloid A (a), haptoglobin (b), tumor necrosis factor alpha (c) and interleukin 2 (d) on days when EEHV1 or EEHV5 was detected in whole blood by PCR. Bars represent the mean ± sem of the prediction from the GLMM; asterisks denote significant differences between categories (** P < 0.01; *** P < 0.001).

**Fig 3:**
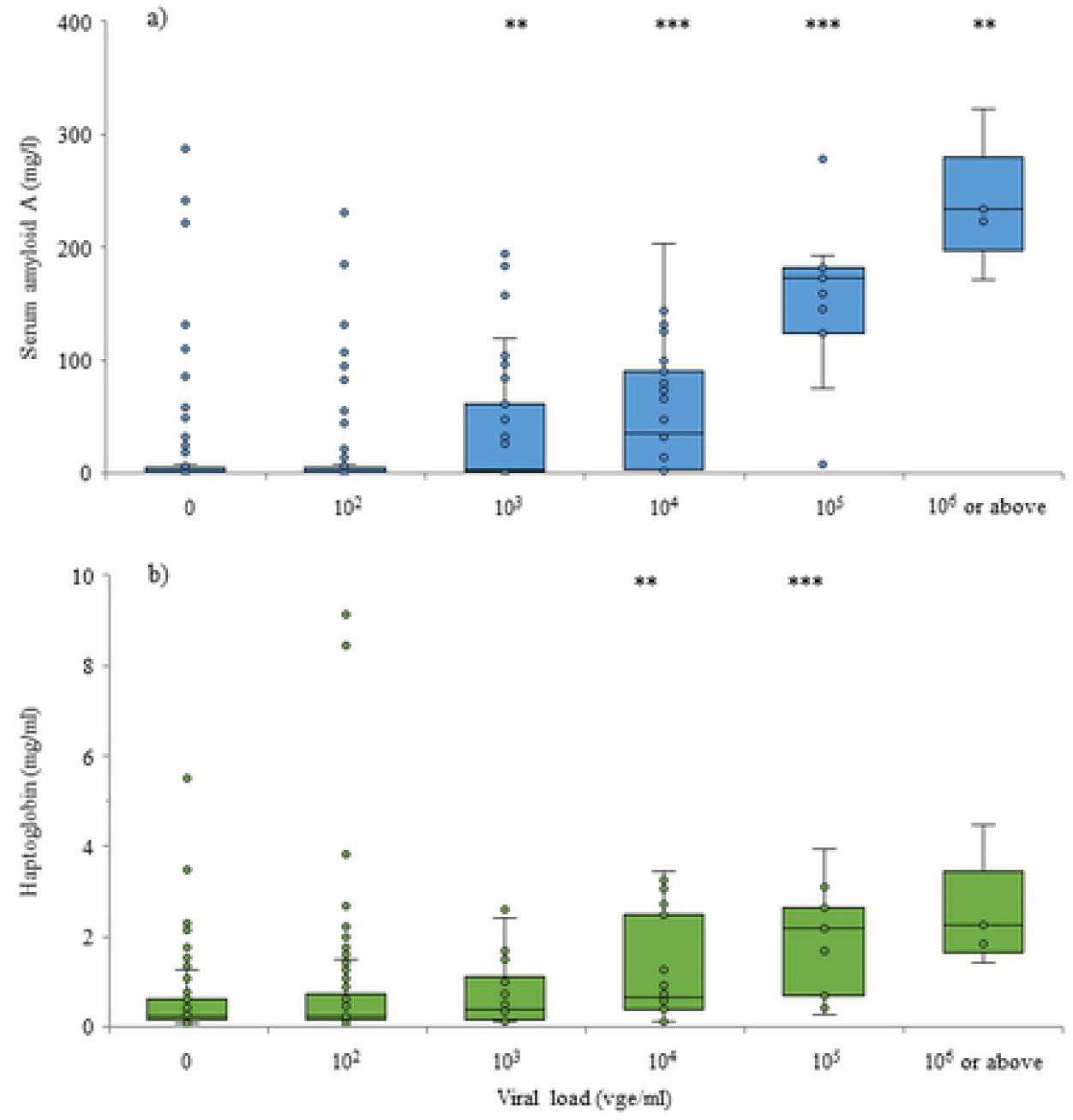
Acute phase protein concentrations with increasing viral load. Serum amyloid A (a) and haptoglobin (b) in elephants with differing EEHV viral loads (EEHV types combined). Box and whisker plots represent the prediction from the GLMM; asterisks denote significant differences compared to 0 vge/ml (** P < 0.01; *** P < 0.001).

Figures 4-6 present longitudinal profiles of serum biomarkers during EEHV1, EEHV5 and EEHV3B viremia, respectively. A female Asian elephant aged 4 years 8 months (Fig 4a) exhibited EEHV1A viremia increasing from 11,000 vge/ml on Day 0 to 500,000 vge/ml by 7 days following initial detection of viral material in whole blood, returning to undetectable levels on Day 25. SAA and HP were both elevated on Day 0 and peaked on Day 1 (203.2 mg/l) and Day 4 (2.7 mg/ml), respectively. SAA then decreased gradually over the next 24 days, whereas HP only remained elevated for 14 days. A second female (Fig 4b) with peak EEHV1A viremia on Day 7 of 10,350 vge/ml showed a similar magnitude increase in SAA (185.7 mg/l; Day 3), but a smaller increase in HP (1.5 mg/ml, Days 9 and 10). Both APPs had decreased to non-viremia levels by Day 21, at which time viremia had reduced to 350 vge/ml. Three females with EEHV5 viremia exhibited lower peak APP concentrations (Fig 5). These three cases with maximum viremia of 23,000 vge/ml, 30,000 vge/ml and 90,000 vge/ml were associated with SAA concentrations below 100 mg/l and HP concentrations below 1.0 mg/ml. In the case presented in Fig 5b, APP concentrations were relatively low during EEHV5 viremia, but elevated SAA and HP concentrations around Day 38 coincided with increased EEHV1 viremia concurrent with EEHV5. Interleukin-2 was below assay detection for all five cases represented in Figures 4 and 5, and TNF-α was only detected in two cases. An 11-year old female with EEHV1 exhibited TNF-α concentrations averaging 32.8 pg/ml that remained fairly consistent throughout the viremia (Fig 4b). Another 11-year-old female with EEHV5 exhibited a steady increase in TNF-α from Day 5 to 14 (Fig 5c); concentrations then remained elevated for the remainder of the sample collection period.

**Fig 4:**
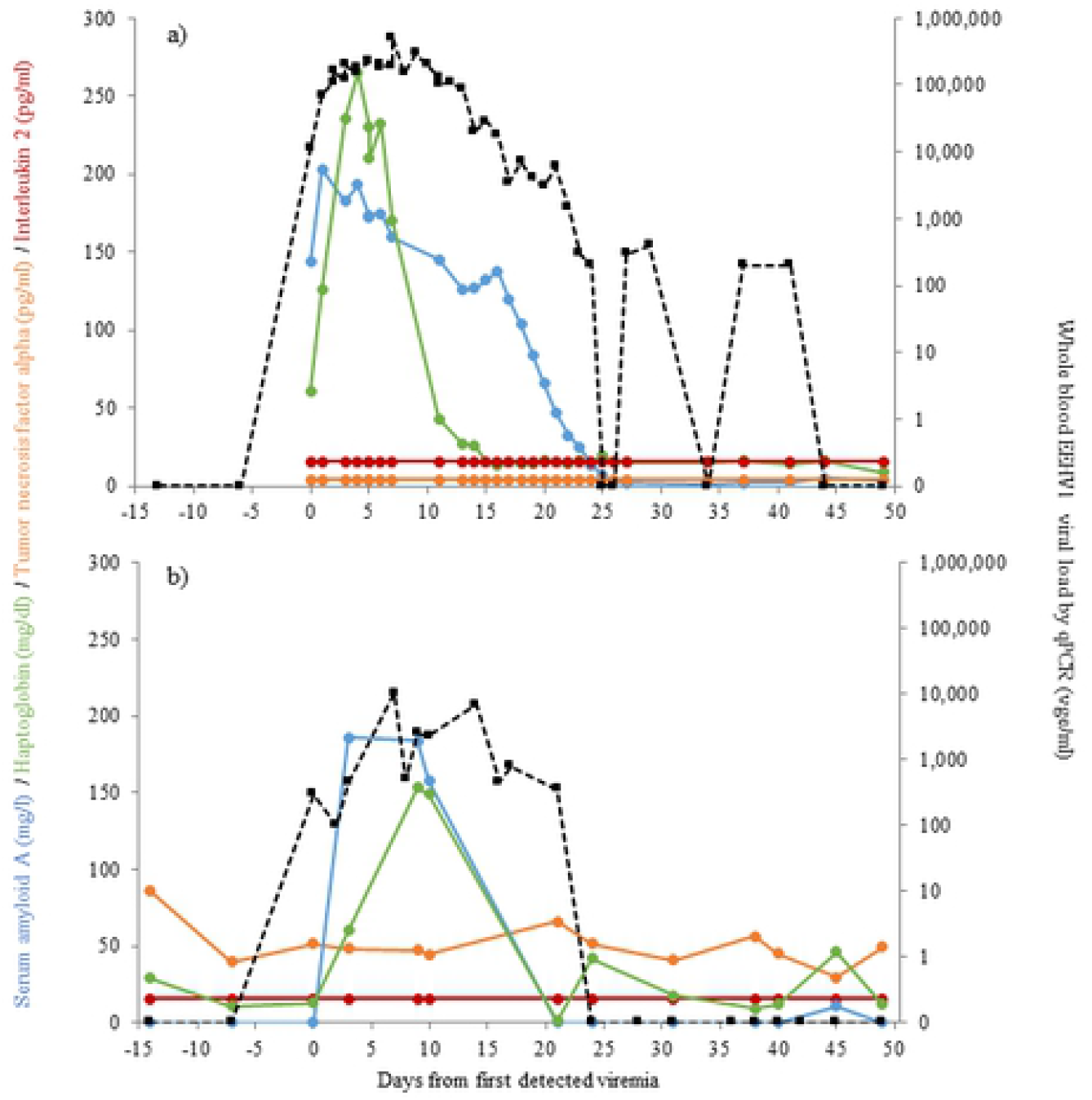
Biomarker concentrations during EEHV1 viremia in two female Asian elephants aged 4 y 8 m (a) and 11 y 11 m (b).

**Fig 5:**
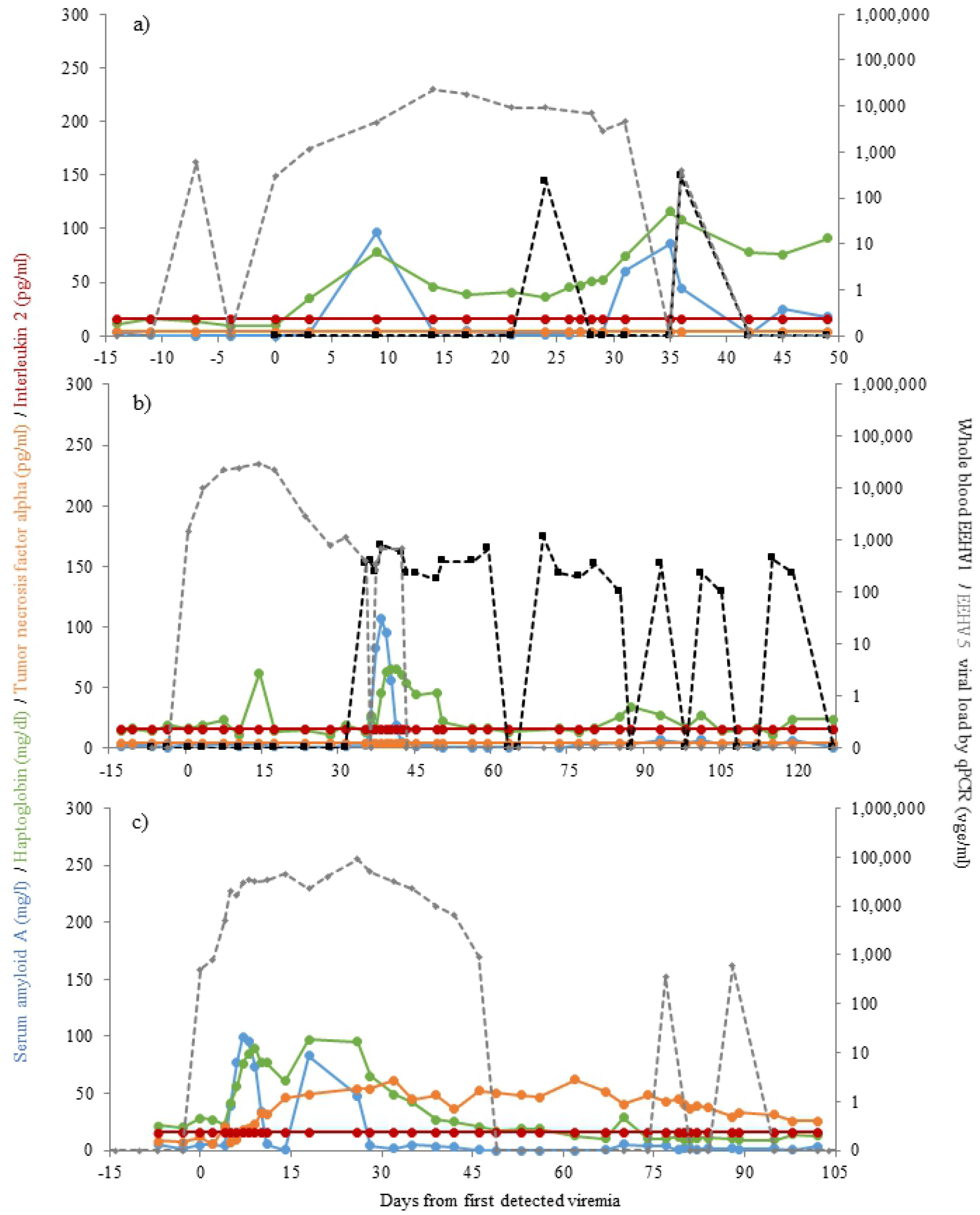
Biomarker concentrations during EEHV5 viremia in three female Asian elephants aged 4 y 10 m (a), 7 y 2 m (b) and 11 y 1 m (c).

**Fig 6:**
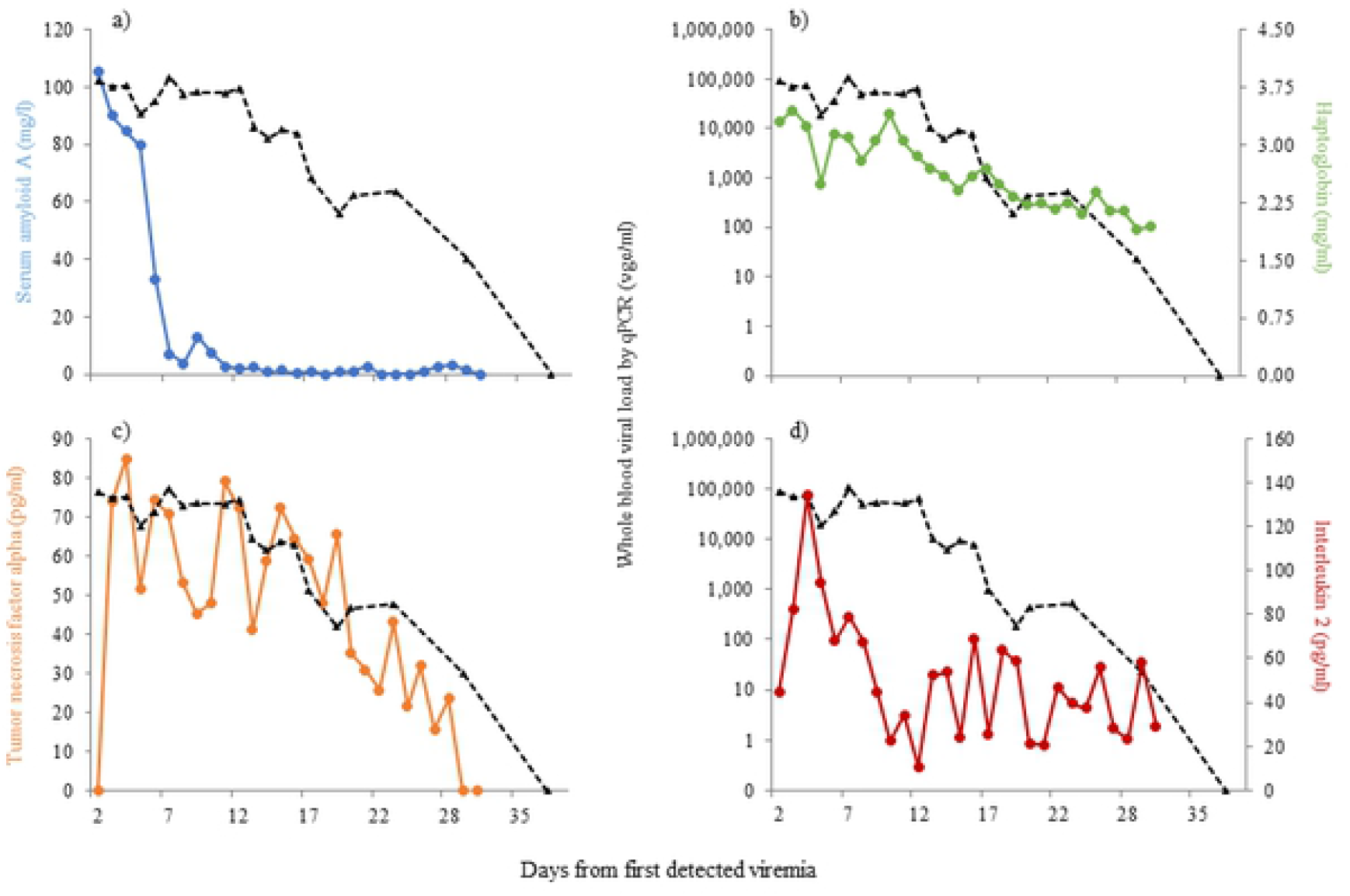
Biomarker concentrations during EEHV3B viremia in a male African elephant aged 4 y 11 m for serum amyloid A (a), haptoglobin (b), tumor necrosis factor alpha (c) and interleukin 2 (d).

The case of EEHV3B in a male African elephant was described by Bronson et al. [30], and here daily samples were additionally analyzed for APP and cytokine concentrations (Fig 6) to provide a longitudinal profile of each biomarker. TNF-α concentrations increased on Day 3 of detected viremia, peaked at 85.1 pg/ml on Day 4 and then gradually declined in parallel with decreasing viremia to below assay detection on Day 30 (Fig 6c). IL-2 was also elevated on Day 3, peaked on Day 4 (134.0 pg/ml) and began to decrease by Day 9, after which concentrations remained relatively stable until the end of the collection period (Fig 6d).

## Discussion

This study measured APP and cytokine biomarkers associated with the immune response during EEHV infection in elephants and found several that could be useful as indicators of disease severity or prognosis. The APPs SAA and HP were both higher in serum from elephants with viremia than those without, and increasing concentrations were associated with increasing viral load. SAA was also higher in cases of EEHV1 compared to EEHV5, and both APPs were higher during fatal cases than in individuals that survived a bout of viremia exceeding 5000 vge/ml. Although concentrations for three (IFN-γ, IL-6 and IL-10) of the five cytokines validated for elephants [45] were undetectable in longitudinal samples, results for TNF-α and IL-2 may show potential as diagnostic tools for studying the immune response to EEHV, although more sensitive techniques would be beneficial.

APPs are useful indicators of inflammation across species [51], and are increasingly used in veterinary medicine [52-54]. As well as being indicative of existing pathology, concentrations can be used as indicators of disease etiology [55, 56], severity [53, 57, 58], and prognosis [54, 59, 60]. A previous study by Stanton and colleagues [29] reported elevated SAA during EEHV1 viremia in Asian elephants, and in samples with greater than 10,000 vge/ml. Our study built upon this by indicating that concentrations of both SAA and HP are impacted by the presence and magnitude of viremia, and correlate with disease outcome. Overall, APPs were elevated in elephants with viral loads of 10^3^ vge/ml or more, and increased with increasing viremia. Visible clinical signs of EEHV HD are often absent until viremia is quite advanced; in the cases reported here viral loads of 19,000 vge/ml EEHV1 and 90,000 vge/ml EEHV were detected by qPCR without accompanying clinical signs of illness. Although clinical signs may not always be apparent, increased APP concentrations indicate that inflammatory processes are underway, and that supportive care should be initiated. We found concentrations of SAA to be greater in EEHV1 infection compared to EEHV5; indeed, APP responses were reduced or even absent in some cases of EEHV5 viremia exceeding 10,000 vge/ml. Historically, EEHV5 has only caused one known death [61] compared to dozens from the two subtypes of EEHV1 [9, 19, 62], suggesting that EEHV5 may be less virulent, and so may be associated with reduced inflammatory responses.

In general, APPs are non-specific, so elevated concentrations may occur in the absence of EEHV viremia. Indeed, some of the samples utilized for this study had been submitted for analysis in response to non-specific clinical signs that upon testing were not associated with EEHV. We recently published reference intervals for several biomarkers in African and Asian elephants and found SAA to be elevated in 83% of active clinical cases, including both infectious and traumatic etiologies, and in 50% of deaths studied [45]. The lack of specificity to EEHV viremia should not preclude the use of APPs as diagnostic tools, however. The rapid and high magnitude increases in SAA in particular are a good indicator of significant inflammatory processes and could be used in combination with other blood parameters, such as lymphocyte and platelet counts [30, 63-66] to ascertain underlying pathology, including EEHV [30, 63-67]. Such data were lacking in this study, so it remains to be determined how they correlate with biomarkers of inflammation. Although changes in HP were of lower magnitude, higher peak concentrations in fatal cases of EEHV HD compared to survivors suggest that this biomarker may also be useful for predicting disease outcomes.

Cytokines have been linked to the pathogenesis of other viral hemorrhagic diseases, such as dengue [68-70], and Ebola [71, 72]. However, what is not yet clear is whether cytokines facilitate the immune response in fighting EEHV infection, or whether over activation contributes to disease progression through a ‘cytokine storm’ that often occurs in the terminal stages of Ebola and other viral diseases [73]. Here we report for the first-time circulating concentrations of cytokines, specifically TNF-α and IL-2, during EEHV viremia. TNF-α is a pro-inflammatory cytokine that plays multiple roles in the immune response, with powerful immunoregulatory, antiviral, cytotoxic, and pro-coagulatory properties [74]. IL-2 supports the growth and differentiation of antigen-activated T lymphocytes, and the development of T-cell immunologic memory. T-cells play an important role in the immune response to viral hemorrhagic disease [75], and are considered to be key in fighting EEHV infection [34]. TNF-α and IL-2 concentrations in this study were low compared to recently published reference intervals [45], and below detection in five individuals. In humans, cytokines are significantly lower in children and increase with age [76]. It is possible, therefore, that our EIAs were of insufficient sensitivity to quantify circulating concentrations of cytokines in the younger calves. However, they did correlate well with EEHV3B viremia in the ∼5 year old African calf; TNF-α concentrations were closely associated with viremia, and IL-2 was elevated during at least Days 3-8 of illness, supporting the proposed involvement of T-cells in EEHV infection [34]. Srivorakul and colleagues [41] recently assessed the expression of cytokines from stimulated PBMCs, and also found increased TNF-α expression in isolated blood cells from an Asian elephant with persistent EEHV4 infection. They proposed that TNF-α may contribute to apoptosis of EEHV-infected cells and to leukocyte migration into affected tissues. Although we cannot determine exact mechanisms, the reduction of TNF-α in parallel with decreasing viremia perhaps supports an association with fighting infection. In other viral hemorrhagic diseases, over-expression of certain cytokines have been associated with damage, including hypotension and hemorrhage [73], and involvement of TNF-α and IL-2 with vascular leakage [75, 77]. We cannot yet rule out such pathological involvement, but our inability to detect cytokine responses in several of the fatal cases of EEHV HD suggests that aberrant over-expression of cytokines may not always be involved in progression of EEHV HD. Indeed, it remains a possibility that an inadequate immune response may be involved in more severe cases of EEHV HD, so more sensitive tools for these and additional cytokines would be beneficial to investigating differential responses between cases, and between African and Asian elephants. To conclude, these results highlight the potential benefit of measuring circulating biomarker concentrations, such as APPs and cytokines, to better our understanding of EEHV viremia and HD.

## Acknowledgments

The authors would like to thank the Albuquerque Biological Park, the Maryland Zoo in Baltimore, the Oklahoma City Zoo, the Oregon Zoo, the Ringling Brothers Center for Elephant Conservation, White Oak Conservation Foundation, and the Saint Louis Zoo for contributing samples and clinical information for use in this study, and Hannah Johnson and Shiling Zhao for laboratory assistance.

